# Cigarette Smoke and E-Cigarette Aerosol Extracts Induce Myelopoiesis and Suppress Inflammatory Cytokine Production

**DOI:** 10.64898/2025.12.21.693601

**Authors:** Jeanette YC Sullivan, Helen X Huang, Jianhong C Heidmann, Francisco Diaz, Camille Brzechffa, Jane H Chen, Nneamaka Iwobi, Irene Hasen, Amanda Ting, Nicole Sparks, Michael T Kleinman, Angela G Fleischman

## Abstract

Tobacco and nicotine use remain the leading preventable drivers of cancer risk, and both direct and secondhand exposure to combustible cigarettes or electronic nicotine devices perturbs immune function and hematopoiesis. Here, we evaluate the impact of e-cigarette vapor and combustible cigarette smoke on *in vitro* cell inflammatory responses and *in vivo* long-term hematopoietic differentiation. In cell-based studies, cigarette smoke extract (CSE) and e-cigarette vapor extract (EVE) consistently suppress LPS-induced TNF-α secretion across macrophage/monocyte models, including primary mouse and human cells and complementary cell lines, indicating a reproducible immunosuppressive effect on mature myeloid cells. Brief ex vivo exposure to CSE also alters myeloid subset composition and modifies the proliferative behavior of Tet2-knockout cells, suggesting that smoke-related cues can reshape competitive dynamics among mutant and wild-type myeloid progenitors. To assess consequences of smoking behavior in vivo, we used a custom nose-cone inhalation system to deliver controlled exposures to combustible cigarette smoke or e-cigarette aerosol to mice. Chronic exposure increased myeloid proliferation, consistent with smoking behavior inducing premature aging of the hematopoietic stem cell pool. Thus, these studies support a model in which tobacco exposures blunt innate immune responsiveness while simultaneously driving myeloid expansion conditions that accelerate hematopoietic aging.

## INTRODUCTION

Combustible cigarette smoke and electronic cigarette aerosol are now two of the most common inhaled exposures that alter immunity and blood formation. Both deliver oxidants and bioactive chemicals that can rewire inflammatory signaling and reshape marrow output. However, their effects are context-dependent, as their impact can vary with dose, timing, and cell type under study. The hematopoietic system contains a wide variety of cells, each with unique physiological functions. At one end of the hierarchy, mature macrophages orchestrate front-line defense and repair; at the other, hematopoietic stem cells (HSCs) integrate inflammatory cues and decide whether to self-renew or differentiate. Tobacco smoke modulates the immune system across the hematopoietic hierarchy, from stem and progenitor cells to mature myeloid populations (Ramanathan et al. 2020), yet side-by-side comparisons of combustible versus e-cigarette exposures across macrophages and HSCs remain limited.

Evidence indicates tobacco smoke exerts bidirectional immune effects; it can provoke inflammation yet also blunt host defense. Smokers show elevated plasma cytokines (Elisia et al. 2020), and a smoking history worsens inflammatory and autoimmune disease courses (Arnson et al. 2010; Mahmoudzadeh et al. 2023). In the lung, smokers have 2-3-fold as many macrophages in their bronchoalveolar lavage fluid (Gonçalves et al. 2011). Paradoxically, smoking also increases the risk of infection (Lugade et al. 2014), likely reflecting impaired adaptive immune responses compromised by chronic inflammation.

Combustible cigarette smoking induces inflammation and oxidative stress and contributes to the pathogenesis of cancers and other diseases; however, the long-term carcinogenic effect of e-cigarettes has yet to be completely recognized. Combustible cigarettes contain tobacco and fillers that are burned and release over 7000 chemicals (Zarcone et al. 2023). E-cigarettes, in contrast, are battery-operated devices that vaporize “e-liquid” containing propylene glycol (PG), vegetable glycerin (VG), flavorings, and nicotine. The accessibility, flavors, and discreet use of e-cigarettes have led to rising nicotine use among adolescents, young adults, and non-smokers. In vitro studies of cigarette smoke extract (CSE), e-cigarette vapor extract (EVE), or their components alone have produced conflicting results, showing both enhanced and repressed cytokine production by monocytes and macrophages. The heterogeneity of e-cigarette formulations further complicates these context-dependent effects. While considerable effort has been devoted to studying the impact of smoking and vaping on mature immune cells, much less is known about how these exposures affect hematopoietic stem and progenitor cells. To address these gaps, we established a standardized in vitro platform to compare the effects of CSE and EVE across multiple macrophage models and to extend these analyses to hematopoietic progenitors.

Cigarette smoke is also strongly associated with the development of clonal hematopoiesis of indeterminate potential (CHIP), the presence of mutant hematopoietic cells without blood count abnormalities (Levin et al. 2022; Dawoud et al. 2020; Kessler et al. 2022; Lee et al. 2024). CHIP occurs when HSCs acquire a mutation that provides a proliferative advantage over their normal counterparts (Marston et al. 2024). The incidence of CHIP increases with age and can be found in about 10% of the population by age 70 (Marston et al. 2024). CHIP increases the risk of blood cancers and cardiovascular diseases (Jakubek et al. 2023; Marnell et al. 2021; Schuermans and Honigberg 2025). Analysis of the UK Biobank (UKB) cohort (n=502,524) identified a significant association (p<0.0005) between smoking behavior and incidence of CHIP (Levin et al. 2022). A separate UKB analysis confirmed this link, showing that myeloid clonal hematopoiesis was particularly enriched among current smokers compared with past smokers (Dawoud et al. 2020). Similarly, exome sequencing from the Geisinger MyCode Community Health Initiative revealed that heavy smoking was strongly associated with CHIP carrier status (Kessler et al. 2022). Importantly, across both the UKB and Geisinger cohorts, CHIP carriers showed an increased risk of hematologic cancers, particularly myeloproliferative neoplasms (Kessler et al. 2022). Together, these findings underscore a consistent epidemiologic link between smoking behavior and aberrant clonal hematopoiesis, but the mechanisms driving this association remain poorly understood.

The most common CHIP mutations occur in the DNMT3A, TET2, ASXL1, and JAK2 genes, which also drive the development of myeloid cancers (Xie and Zeidan 2023). These CHIP clones emerge because they gain a competitive advantage under hematopoietic stress and aging. At the stem and progenitor level, inflammatory and oxidative stress cues transiently push HSCs into cycle and bias lineage output toward the myeloid fate; chronic or repeated stress can erode self-renewal and remodel progenitor composition (Pietras 2017). TET2, an epigenetic regulator of DNA methylation, is the second most frequently mutated CHIP gene (Moran-Crusio et al. 2011; Abdel-Wahab et al. 2012; Cobo et al. 2022). Somatic *TET2* mutations can be found in 50% of chronic myelomonocytic leukemia, 30% of patients with myelodysplastic syndromes or myeloproliferative neoplasms, and are associated with worse outcomes in acute myeloid leukemia (AML) (Moran-Crusio et al. 2011). The transformation of underlying CHIP mutations into official malignancies is a key clinical concern, but the factors that control their emergence remain poorly understood. Here, we leverage the associations between CHIP, inflammation, and smoking to investigate how these factors contribute to mutant TET2 hematopoiesis.

## MATERIALS AND METHODS

### Mice

All animal procedures were performed under the approval of the Institutional Animal Care and Use Committee at the University of California, Irvine. Wildtype C57BL/6J (Stock No. 000664) and Tet2^-/-^ mice (Stock No. 023359) were purchased from The Jackson Laboratory (Bar Harbor, ME) at 8- to 10-weeks old and housed in specific pathogen-free facilities on a 12-hour light/dark cycle. Euthanasia was performed using an overdose of isoflurane, and euthanasia was confirmed by cervical dislocation. Where indicated, complete blood counts were obtained using the automated cell counter machine (ABCVet Hemalyzer, scilVet).

### Isolation of human peripheral blood mononuclear cells (PBMCs)

Peripheral blood was collected from healthy blood donors in EDTA tubes. Peripheral blood mononuclear cells (PBMCs) were isolated using the Lymphoprep^TM^ density gradient medium (Stemcell).

### THP-1 and RAW264.7 cells

RAW 264.7 cell lines were obtained commercially. Cells were revived from a previously frozen third passage stock and cultured in RPMI 1640 medium supplemented with 10% fetal bovine serum (FBS) and 1% penicillin/streptomycin (R10) in 10 mm tissue culture-treated Petri dishes. Cultures were maintained at 37°C in a 5% CO₂ incubator. Once the cells reached confluency, they were passaged using 1 mL of trypsin for 5 minutes to facilitate detachment. Cells were gently scraped and transferred to a 50 mL conical tube containing 40 mL of R10. From this, 10 mL of the cell suspension was replated into a fresh 10 mm Petri dish for continued culture. The remaining cells were seeded into 96-well tissue culture-treated plates and allowed to adhere overnight in the incubator before stimulation. Cells were continuously passaged from passage 3 through passage 15.

THP-1 cell lines were obtained commercially (American Type Culture Collection, USA). Cells were thawed from a previously frozen stock and cultured in RPMI 1640 medium supplemented with 10% fetal bovine serum (FBS) and 1% penicillin/streptomycin (R10) in T-25 flasks. Cells were maintained at 37°C in a 5% CO₂ incubator. Cells were continually passaged from passage 3 through 15.

### Bone marrow-derived macrophages (BMDMs)

Bone marrow was extracted by flushing the femurs and tibias with 10 mL of staining buffer using a syringe through a 45 µm mesh strainer. The collected marrow was subjected to red blood cell lysis using ACK buffer on ice for 20 minutes, followed by washing and centrifugation. The resulting cell pellet was resuspended in DMEM supplemented with 10% fetal bovine serum (FBS) and 1% penicillin/streptomycin (D10). A small aliquot of the suspension was analyzed by flow cytometry to determine the total bone marrow cell count. To generate bone marrow-derived macrophages (BMDMs), 20 million bone marrow cells were resuspended in 10 mL of D10 supplemented with 30% L929 cell-conditioned media. Cells were seeded in non-tissue culture-treated 10 mm Petri dishes and incubated at 37°C in a 5% CO₂ atmosphere. After 3 days, each dish was supplemented with an additional 3 mL of fresh D10 + 30% L929 media. On day 5, media in each dish was replaced entirely with 10mL fresh D10 + 30% L929 media. After 7 days, the supernatant was discarded, and the macrophages were incubated with 1 mL of trypsin for 5 minutes. Macrophages were detached by scraping gently and collecting into D10.

### Cigarette smoke extract (CSE) preparation

Cigarette smoke extract (CSE) was prepared from mainstream cigarette smoke using a standard protocol. Briefly, mainstream smoke from one 2R4F research cigarette was drawn into serum-free media by continuous negative pressure and filter-sterilized to obtain 100% CSE. Volume/volume percentage (v/v %) dilutions were used for in vitro exposures.

### E-cigarette vape extract (EVE) preparation

E-cigarette smoke extract (EVE) was prepared using the Naked brand Melon flavored e-liquid (65:35 VG:PG with 6mg nicotine) in a Vaporesso XROS 3 Pod System. The vape liquid was purchased from a local shop in Irvine, CA. The Vaporesso device was attached to an impinger, which is attached to the vacuum port of a tissue culture hood. To mimic inhalation, the vacuum was turned on for 4.3 seconds every minute for a duration of 75 minutes, drawing the e-cigarette vapor into 25mL of media. This protocol was developed based on the collection methods described in Williams et. al. (Williams et al. 2019).

### Nose-only inhalation exposures

Exposures were performed at the Air Pollution Health Effects Laboratory at the University of California, Irvine. Exposures occurred 2 hours per day, 4 days/week, for 8 weeks. Between exposures, the mice were housed 4 per cage in an atmosphere-controlled room on a 12-hr light/dark cycle in an Association for the Assessment and Accreditation of Laboratory Animal Care (AAALAC) accredited animal housing facility at the University of California, Irvine vivarium. Aerosols were generated using a 2-s puff with a 60-s interval between puffs (35 mL puff volume) based on the ISO standard cigarette puff protocol (ISO 3308:2012) using a custom-built smoking system (Figure 9B). This system uses a peristaltic pump to draw in aerosolized smoke from a lit combustion cigarette (1R6F Certified Reference Cigarette, Center of Tobacco Reference Products (CTRP), University of Kentucky) and directs it into a nose-only exposure manifold (In-Tox Products, Moriarty, NM, USA). For the e-cigarette exposure, the peristaltic pump draws in aerosolized vapor from a Nichrome 70-watt E-cigarette. The e-liquid used was made by mixing equal volumes of propylene glycol (Millipore, Burlington, MA) and vegetable glycerin (Fisher Chemical, Hampton, NH) with 1.5% of nicotine (ThermoFisher, Waltham, MA). Control mice were exposed to air purified over potassium permanganate-impregnated alumina beads, activated carbon, and high-efficiency particulate air (HEPA) filters, concurrently with the combustion cigarette or e-cigarette exposures.

*The following information was previously published and is provided here for convenience* (Ramanathan et al. 2023). The nose-cone inhalation exposure system is advantageous over whole-body exposure systems because it minimizes fur contamination and potential non-relevant exposure due to routine grooming that occurs in a whole-body exposure regime. During exposures, animals were held in individual exposure tubes that were connected to the exposure manifold and positioned with just the snout exposed to the exposure atmosphere. Exhaust ports surrounding each nose cone were under slight negative pressure to direct the flow of fresh aerosols to the animal’s breathing zone and exhaust exhaled air, thereby minimizing the potential for re-breathing secondary vapor and preventing CO_2_ buildup.

### Bone marrow transplantation

Fresh bone marrow was harvested from euthanized adult mice. Recipient mice were lethally irradiated with a single dose of 8Gy and transplanted retro-orbitally with approximately 1.2×10^6^ whole bone marrow cells. For the competitive wild-type (WT) and Tet2^-/-^ transplant, mice were transplanted with a ratio of 93% WT to 7% Tet2^-/-^ cells.

### Platelet counting

Platelet counting was performed as adapted from Masters & Harrison, 2014 (Masters and Harrison 2014). Peripheral blood from the saphenous vein was mixed 1:1 with EDTA, diluted in staining buffer with 0.2 microliters of FITC CD61, and incubated for 20 minutes. Samples were further diluted and run on a Novocyte 3000 flow cytometer (Agilent Technologies) to obtain absolute platelet counts.

### Flow cytometry of mouse peripheral blood/bone marrow

To determine peripheral blood cell frequencies and chimerism, red blood cells were lysed with ammonium-chloride-potassium (ACK) buffer. Cells were stained with anti-mouse antibodies specific to CD45.1, CD45.2, CD11b, CD19, CD11b, Ly6G, CD3, CD4, CD8, and Ly6C. For the immature stem and progenitor populations, bone marrow was stained with anti-mouse antibodies CD34, CD150, CD16/32, CD48, cKit (CD117), lineage cocktail (containing: CD3, Ly-6C, CD11b, CD45R, Ter-119), and Sca-1. All antibodies were used at a 1:500 dilution factor as recommended.

### Colony formation assay

Colony formation assays were performed as indicated by StemCell Technology protocols. Whole bone marrow cells from WT and Tet2^-/-^ were incubated with increasing concentrations of cigarette smoke extract (CSE) or e-cigarette vape extract (EVE) overnight. Cells were washed thoroughly and plated at a concentration of 10,000 cells/ml in Methocult medium (M3224, StemCell) with IL-3 at 10ng/mL and SCF at 50ng/mL. Colonies were counted 7-12 days later.

### Measurement of cytokine secretion

Concentrations of TNF-α were determined in conditioned media supernatant from cells incubated with 10ng/mL lipopolysaccharide (LPS, Sigma) and increasing concentrations of CSE and EVE for 24 hours. Samples were frozen at −80°C until analysis. Commercial enzyme-linked immunosorbent assay (ELISA) kits for human TNF-α (88-7346-88, Invitrogen/ThermoFisher) and mouse TNF-α (88-7324-88, Invitrogen/ThermoFisher) and used in accordance with the manufacturer’s instructions.

### Statistical analysis

Data are presented as means ± standard error of the mean (SEM). GraphPad Prism (GraphPad Software, San Diego, CA) was used for all statistical analyses. Differences between mean values within groups were determined using ANOVA or unpaired t-tests where indicated.

## RESULTS

### Standardized in vitro model reveals differential inhibitory effects of cigarette smoke and e-cigarette exposure on macrophage TNF-α responses

To investigate how smoke or e-cigarette exposure influences inflammatory responses, we established a standardized workflow in which all cell culture and analytic protocols were consistent across experiments. Single-use aliquots of the original batches of CSE and EVE, ensuring that the only variable was the cell type under study. This approach enabled a direct, side-by-side comparison of different monocyte and macrophage models. Because higher concentrations of CSE and EVE can impair cell proliferation and viability, we selected minimal concentrations (0.5-2%), which did not induce significant cell death under our culture conditions.

We first examined how CSE and EVE influence tumor necrosis factor-alpha (TNF-α) production in response to lipopolysaccharide (LPS). Cells were exposed to CSE and/or EVE concurrently with LPS stimulation, and supernatant was collected 24 hours later to quantify TNF-α production by ELISA. To broadly evaluate these effects, we tested both human and murine macrophage cell lines, as well as primary macrophages derived from human peripheral blood monocytes and mouse bone marrow (Figure 1). In RAW264.7 cells, a murine macrophage cell line widely used in immunology and toxicology studies, CSE significantly suppressed TNF-α secretion in a dose-dependent manner (Figure 1A). Suppression of TNF-α by CSE was also confirmed using intracellular cytokine staining (Supplemental Figure 1). In contrast, EVE prepared from a commercially available melon-flavored e-liquid did not significantly alter TNF-α levels (Figure 1A).

**Figure 1.**
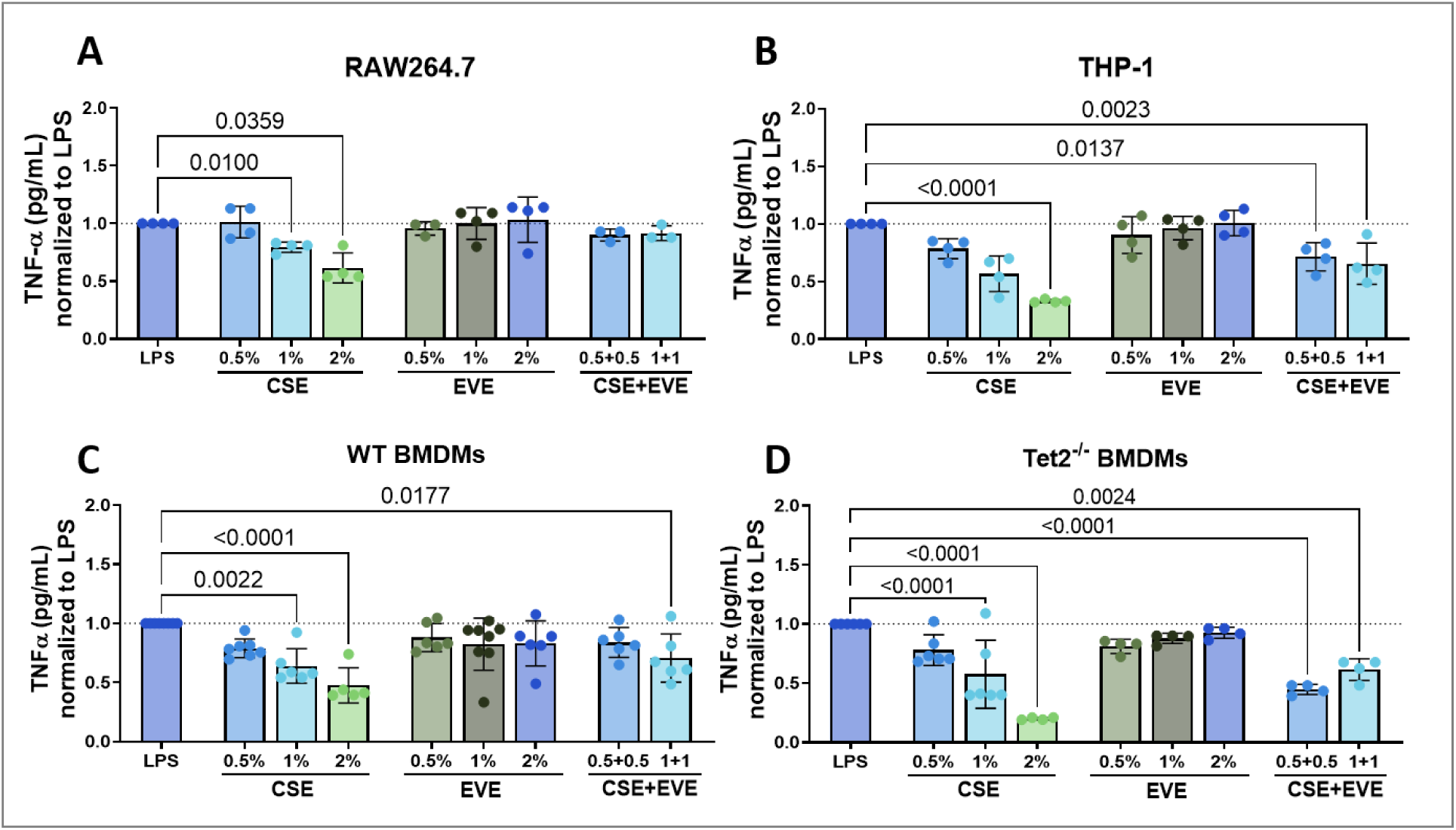
Cigarette smoke inhibits LPS-induced TNF-α secretion in multiple macrophage models. Adherent macrophages were washed and stimulated with 10ng/mL LPS and 0-2% CSE, 0-2% EVE, or a combination of CSE+EVE. Supernatant was collected at 24H and frozen at −80 °C until analyzed by TNF-α ELISA. Ratio of TNF-α is calculated by normalizing the amount of TNF-α secreted with CSE or EVE by the amount of TNF-α secreted by the LPS-stimulated cells alone. (A) RAW264.7 macrophages were washed, plated in 96-well plates, and incubated overnight for adherence pre-stimulation. Results are the ratios of 4 independent experiments with n=4-10/group/experiment. (B) THP-1 monocytes were plated in 96-well plates and incubated with PMA for 48 hours to induce macrophage differentiation and washed before stimulation. Results are the ratios of 4 independent experiments with n=4-10/group/experiment. (C) Bone marrow was freshly isolated from wild-type and (D) Tet2-deficient mice and incubated for 7 days with 15% L929-conditioned media (to provide mM-CSF) on non-tissue culture treated petri dishes to induce differentiation of bone-marrow-derived macrophages (BMDM). BMDMs were washed and replated prior to LPS/CSE/EVE exposure. Results are the ratios of 5-8 independent experiments with n=4-8 replicates/group/experiment. Significance is calculated by one-way ANOVA. Values represent means ± SEM.

**Figure 2.**
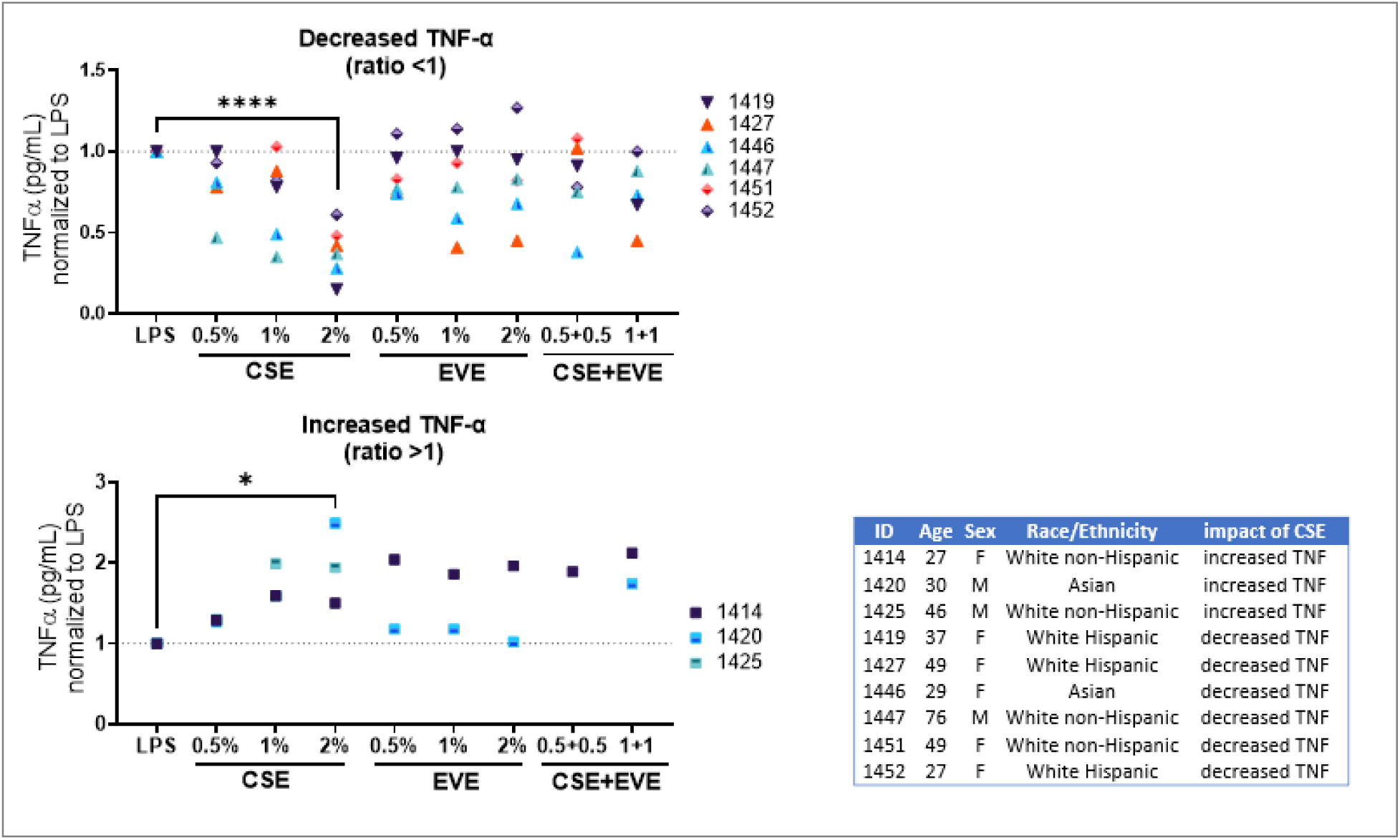
Human PBMCs are diverse and display either a “stimulatory” or “inhibitory” response. Normal human peripheral blood mononuclear cells were exposed to 10ng/mL LPS and 0-2% CSE, 0-2% EVE, or a combination of CSE+EVE. Supernatant was collected at 24H and frozen at −80 °C until analyzed by TNF-α ELISA. Ratio of TNF-α is calculated by normalizing the amount of TNF-α secreted with CSE or EVE by the amount of TNF-α secreted by the LPS-stimulated cells alone. Results are the mean of n=4-8 replicates per donor. Significance is calculated by one-way ANOVA. Values represent means ± SEM.

We then applied the in vitro smoke and e-cigarette exposure protocol to THP-1-derived macrophages, a human monocytic cell line frequently used in immunology and oncology research. THP-1 monocytes were differentiated into macrophages with Phorbol 12-myristate 13-acetate (PMA) before stimulation. Like RAW264.7 cells, THP-1 macrophages showed a dose-dependent reduction in TNF-α secretion in response to CSE (Figure 1B). While EVE alone did not significantly suppress TNF-α production, co-exposure with CSE further reduced secretion compared with CSE alone (Figure 1B).

We next tested how CSE and EVE affect primary bone marrow-derived macrophages (BMDMs). To assess genotype-specific responses, we generated BMDMs from wild-type C57BL/6 mice and from Tet2^⁻/⁻^ mice, given that clonal hematopoiesis, most commonly involving TET2 mutations, is enriched in smokers. This design enabled us to ask whether CHIP-mutant macrophages respond differently to smoke exposure than their wild-type counterparts. Like the cell line macrophages, we observed a dose-dependent inhibition of TNF-α in both WT and Tet2^-/-^ BMDM (Figure 1C, D). When directly comparing WT vs Tet2^-/-^ macrophages, the impact of smoke was greater in Tet2^-/-^ macrophages, specifically at 2% CSE and 0.5+0.5 CSE + EVE, demonstrating a differential effect of smoking on Tet2^-/-^ macrophages. These findings indicate that when a standardized protocol is used, primary macrophages and immortalized cell lines respond similarly to CSE and EVE, both demonstrating suppressive effects of CSE.

We next applied this workflow to human peripheral blood mononuclear cells (PBMCs) freshly isolated from whole blood of healthy donors. Altogether, the responses to CSE and EVE were variable across individuals, without a uniform pattern of TNF-α inhibition or stimulation overall. However, when analyzed on a per-donor basis, the data revealed two distinct response groups: an anti-inflammatory group and a pro-inflammatory group. These groups were not associated with age, sex, or medication use, and smoking history was unavailable. Most donors (6 of 9) exhibited reduced TNF-α secretion after CSE exposure, with some also showing suppression at higher EVE doses (TNF-α ratio <1). In contrast, three donors demonstrated the opposite effect, with enhanced TNF-α secretion following exposure to CSE or EVE. These findings underscore the heterogeneity of human immune responses to smoke- and vape-derived exposures, which may reflect variability in trained immunity that shapes innate immune responsiveness.

### Cigarette smoke and e-cigarette extracts suppress hematopoietic progenitor function with partial resistance in Tet2-deficient cells

To examine how CSE and EVE influence hematopoietic progenitor function, we utilized methylcellulose colony formation assays to quantify myeloid progenitor activity. Given that smoking increases the risk of clonal hematopoiesis of indeterminate potential (CHIP), we tested whether progenitors carrying a common CHIP mutation, Tet2 loss-of-function, respond differently to these exposures. Whole bone marrow from wild-type or Tet2^⁻/⁻^ mice was incubated overnight with CSE, EVE, or their combination, washed, and plated in methylcellulose. Myeloid colonies were enumerated after 7–10 days. Both CSE and EVE significantly reduced colony formation in wild-type and Tet2^⁻/⁻^ cells (Figure 3). However, Tet2^⁻/⁻^ progenitors retained higher clonogenic potential compared with wild-type cells, particularly at 1% and 2% CSE concentrations (Figure 3). These results suggest that Tet2 deficiency confers relative resistance to the suppressive effects of cigarette smoke on hematopoietic progenitors.

**Figure 3.**
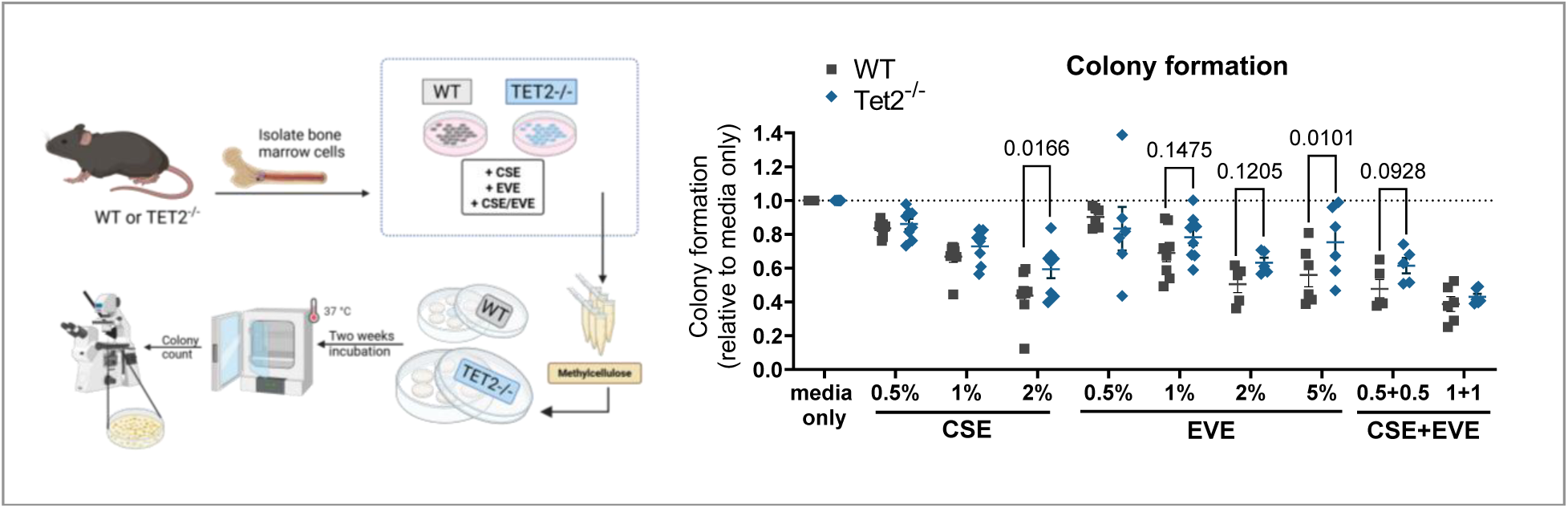
Cigarette smoke and e-cigarette extracts suppress hematopoietic progenitor function with partial resistance in Tet2-deficient cells. Wildtype and Tet2^-/-^ mouse bone marrow cells were incubated overnight with increasing concentrations of cigarette smoke extract and/or e-cigarette vape extract, washed twice, then plated in methylcellulose supplemented with SCF, IL-3, and EPO. Colonies were enumerated 7-10 days later. Significance is calculated by one-way ANOVA. Values represent means ± SEM.

### Evaluating the impact of CSE and EVE on the clonal competition of Tet2-deficient cells

Based on the observed effects of overnight CSE and EVE exposure on hematopoietic stem and progenitor cell function in vitro, we next investigated how this exposure impacts in vivo hematopoiesis. Given the relative resistance of Tet2^⁻/⁻^ cells in vitro, we also asked whether smoke or e-cigarette exposure might promote their expansion or alter their phenotypic behavior in vivo. To test this, we performed competitive bone marrow transplants using cells pre-exposed to CSE, EVE, or the combination (Figure 4). Whole bone marrow cells from wild-type (CD45.1) and Tet2^-/-^ (CD45.1/2) mice were mixed at a ratio of 93:7 and cultured overnight in either R10 media only (RPMI + 10% FBS) or R10 with 0.5% CSE, 1% EVE, or 0.5% CSE and 1% EVE (Figure 4). An aliquot of cells collected after culture confirmed that the input ratio of 7% Tet2^⁻/⁻^ to 93% WT was preserved across all conditions. These co-cultured cells were then transplanted into lethally irradiated CD45.2 WT recipients (Figure 4).

**Figure 4.**
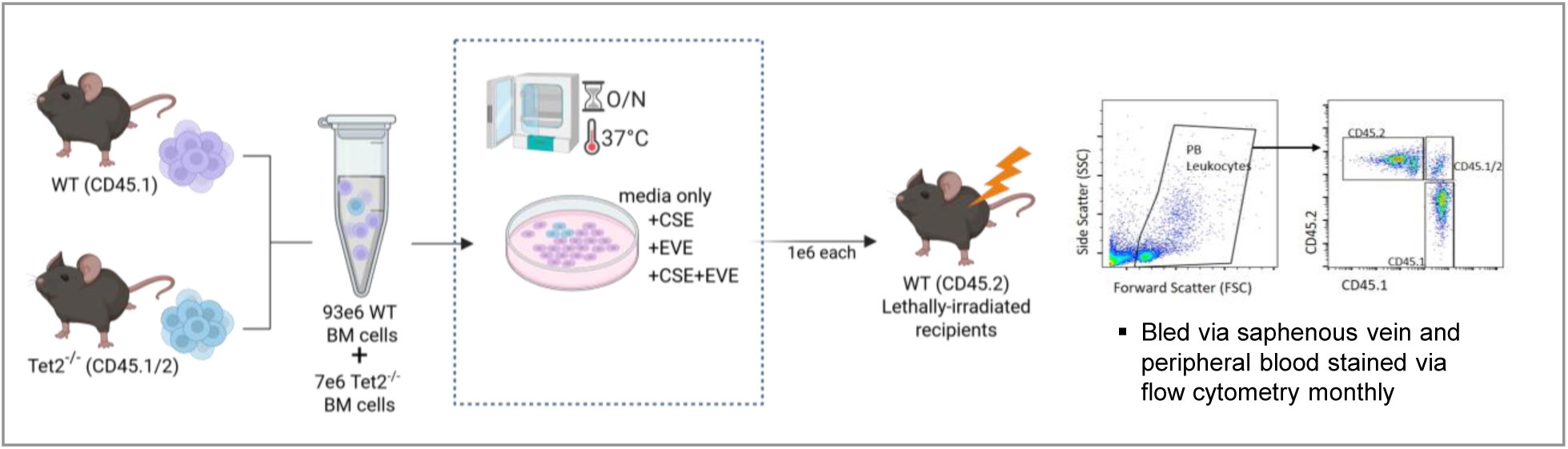
Experimental setup of WT: Tet2-/- competitive transplant with bone marrow co-cultured overnight in smoke or e-cigarette extract. Whole bone marrow was freshly isolated from WT (CD45.1) and Tet2-/- (CD45.1/2) mice and mixed at a ratio of 7% Tet2-knockout to 93% WT. The mixed cells were exposed to CSE or EVE overnight. After incubation, the cells from each were confirmed to still have 7% Tet2-/- and 93% WT by flow cytometry staining (representative CD45.1, CD45.2, and CD45.1/2 gating shown). The cells were washed and transplanted into lethally irradiated CD45.2 WT mice, n=9-10 per group. Mice were bled via the saphenous vein monthly to determine peripheral blood chimerism and composition.

### A single exposure to CSE or EVE reprograms HSCs toward myeloid skewing

We followed peripheral blood counts monthly following transplantation using flow cytometry (Figure 4). In the mice transplanted with cells cultured in media only, myeloid cell frequencies remained stable over time (Figure 5A). In contrast, mice transplanted with CSE and EVE-exposed BM showed a significant (p < 0.05) increase in the percentage of myeloid cells over time (Figure 5A). This myeloid bias, a hallmark of HSC aging (Rossi et al. 2007), suggests that transient exposure to cigarette smoke or e-cigarette vapor can alter hematopoietic function in vivo. We further investigated specific myeloid subsets, focusing on neutrophils and monocyte/macrophages. E-cigarette vapor has been shown to impair neutrophil functions, including chemotaxis, phagocytosis, and NETosis (Corriden et al. 2020; Jasper et al. 2024). Non-exposed bone marrow cultured in media-only did not significantly increase the post-transplant proportions of neutrophils or monocytes and macrophages over time (Figure 5B). However, exposure to e-cigarette extract significantly increased the number of neutrophils in the peripheral blood (Figure 5B). In contrast, exposure to CSE or the combination of CSE+EVE increased the proliferation of monocytes and macrophages (Figure 5C). CSE also modestly increased the absolute platelet count (Figure 5D), consistent with a chronic inflammatory state. The number of lymphoid B and T cells in the peripheral blood was overall unaffected by pre-incubation with CSE or EVE. Together, these findings show that even a single pre-transplant exposure to cigarette smoke or e-cigarette extract can durably reprogram bone marrow, biasing hematopoiesis toward myelopoiesis, a hallmark feature of hematopoietic aging.

**Figure 5.**
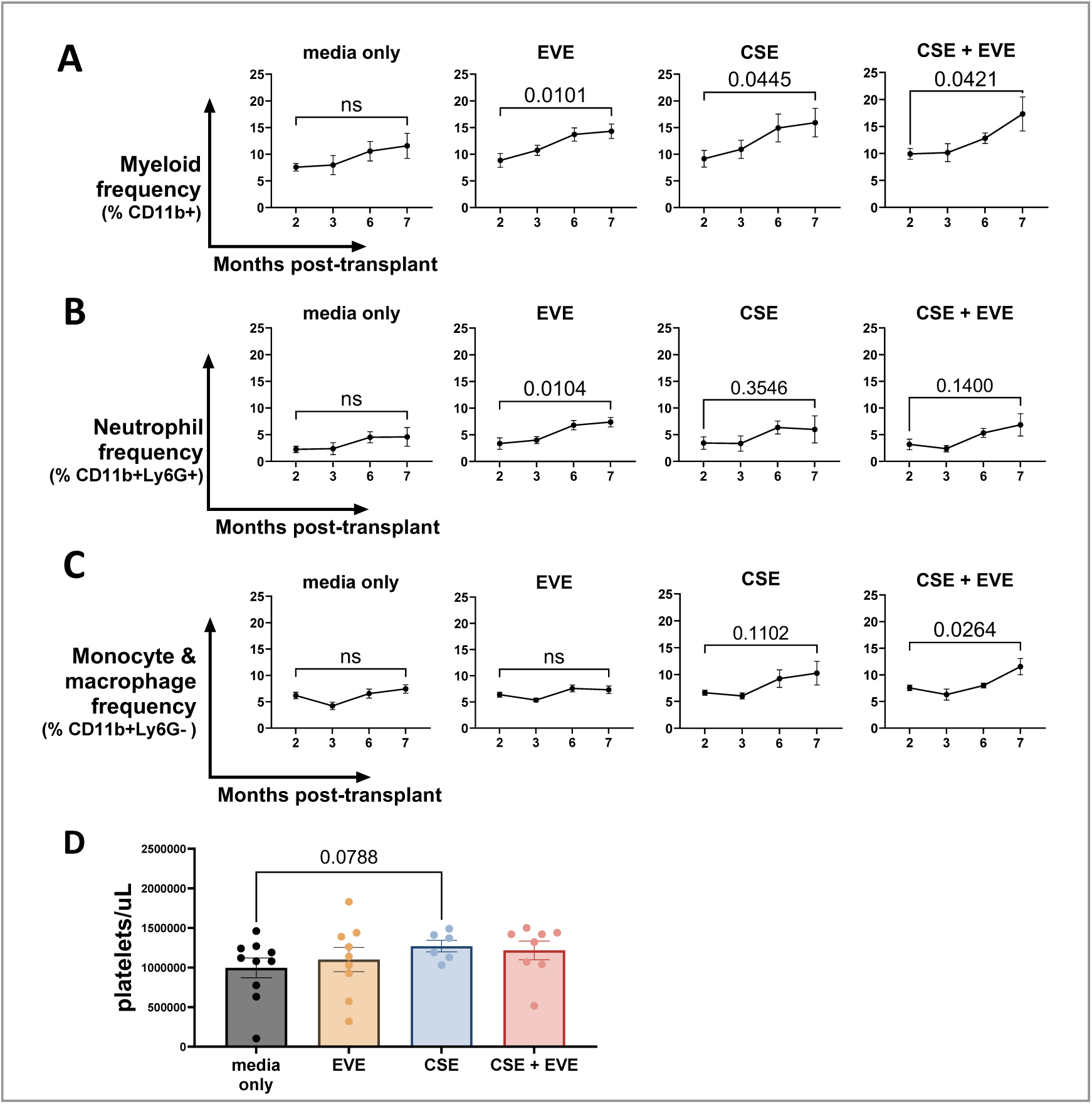
Exposure to CSE or EVE in HSCs induces long-term increases in circulating myeloid cell populations. Flow cytometry was used to analyze the peripheral blood myeloid and lymphoid composition of chimeric mice transplanted with CSE- or EVE-exposed cells. Changes in the frequency of (A) CD11b+CD19- myeloid cells, (B) CD11b+Ly6G+ neutrophils, and (C) CD11b+Ly6G- monocytes of all CD45+ leukocytes in the peripheral blood over time. (D) Platelet counts in the peripheral blood at month 7. Values represent means ± SEM. Significance calculated by unpaired t-tests (A-C) and one-way ANOVA (D).

### EVE promotes the expansion of Tet2-deficient mutant cells in the peripheral blood

We next asked whether overnight exposure to CSE or EVE would alter the selective advantage of Tet2-deficient cells in vivo. Based on our in vitro methylcellulose data, we anticipated that smoke extract might preferentially suppress wild-type progenitors, thereby enhancing Tet2^⁻/⁻^ expansion after transplant. To test this, we measured peripheral blood chimerism monthly after competitive hematopoietic stem cell transplantation. As expected, Tet2^⁻/⁻^ cells expanded in all mice, rising from the 7% input proportion to about 30% of donor-derived cells by one month post-transplant and about 60% by two months (Figure 6A). At this point, the expansion of Tet2^⁻/⁻^ cells begins to diverge between the groups; Tet2^⁻/⁻^ continued to proliferate in recipients of cells cultured in EVE, ultimately reaching about 80% of donor-derived cells (Figure 6A). In mice transplanted with non-exposed or CSE-exposed bone marrow, Tet2^⁻/⁻^ expansion plateaued or fell slightly to around 50% (Figure 6A). These results suggest that e-cigarette aerosols provide a proliferative advantage for the clonal expansion of Tet2-deficient cells.

**Figure 6.**
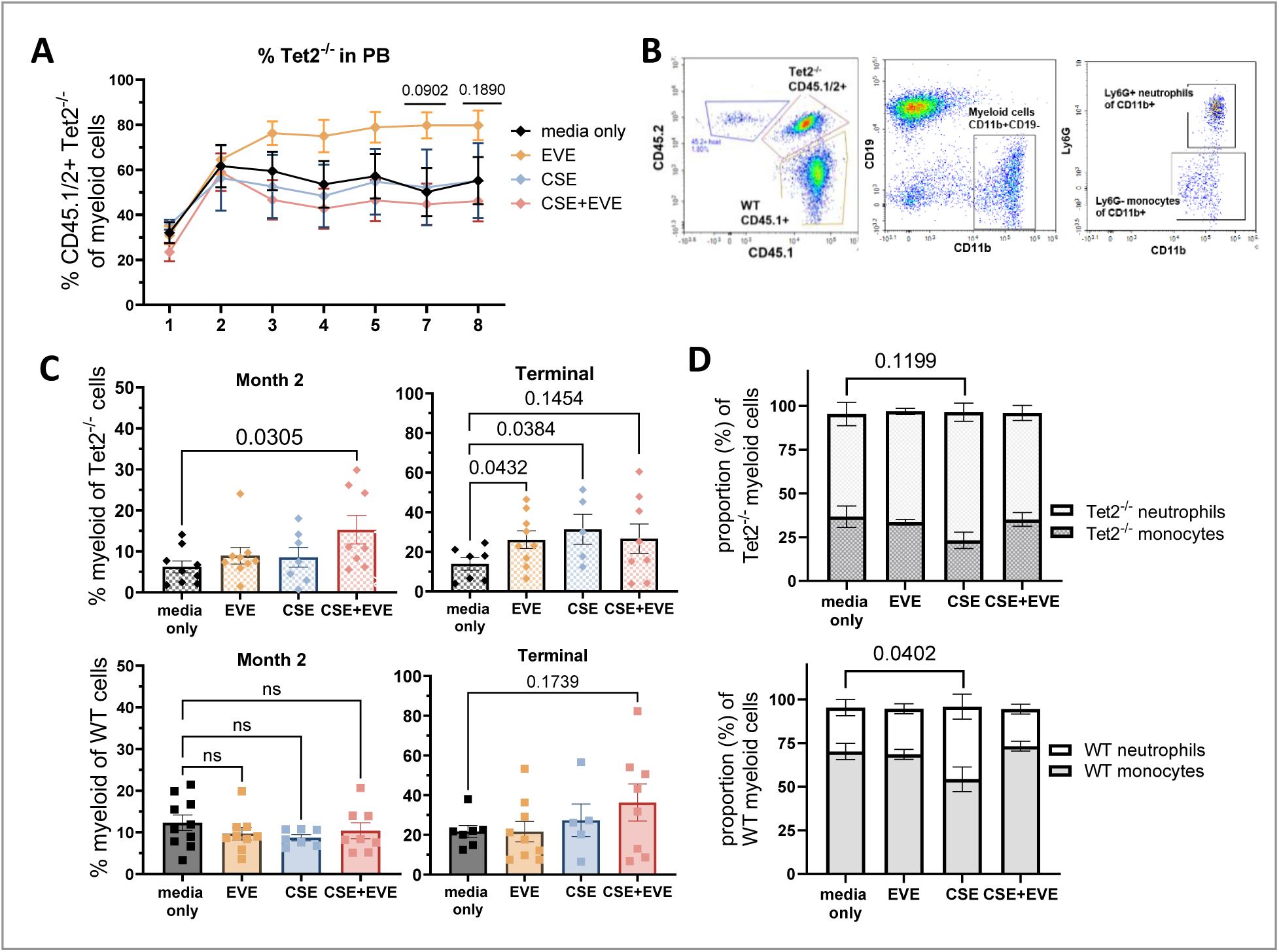
EVE exposure promotes the expansion of Tet2^-/-^ cells within the peripheral blood in a competitive transplant. (A) Frequency of donor-derived CD45.1/2+ cells of myeloid CD11b+CD19-CD45+ leukocytes in the peripheral blood as measured by flow cytometry gating described in (B). P-values shown are the one-way ANOVA results between the EVE and the non-exposed media-only group. (C) The percentage of CD11b+CD19- myeloid cells in Tet2^-/-^ (top) and WT (bottom) mice at months 2 (left) and month 10 at termination (right). (D) The proportions of Ly6G+ neutrophils and Ly6G- monocytes within the CD11b+CD19- myeloid populations of the CD45.1/2+ Tet2^-/-^ and CD45.1 WT leukocytes. P-values shown are the results of one-way ANOVA analyses of the Ly6G+ neutrophil frequencies. Values represent means ± SEM.

To analyze the impacts of CSE or EVE exposure on each of these WT or Tet2^⁻/⁻^ populations individually, we used flow cytometry to gate on the Tet2^⁻/⁻^ CD45.1/2+ and WT CD45.1+ cells of all peripheral blood CD45+ leukocytes. We then quantified the frequency of CD11b+ myeloid cells, CD11b+Ly6G+ neutrophils, and CD11b+Ly6G- monocytes (Figure 6B). Co-exposure to CSE + EVE significantly increased the proportion of myeloid cells within the Tet2^⁻/⁻^ compartment (CD45.1/2⁺) after 2 months (Figure 6B). At termination (10 months), the EVE- and CSE-exposed Tet2^⁻/⁻^ cells had significantly increased myeloid cell proportions (Figure 6B). These dynamics indicate that both CSE and EVE separately increase myelopoiesis of mutant Tet2^-/-^ cells, and the combination of the two accelerates this impact. In contrast, wild-type cells showed no significant changes in overall myeloid CD11b+ frequencies across conditions through month 2 to termination (Figure 6B).

Closer examination of myeloid cells revealed that although the overall frequency of myeloid cells was unchanged, CSE altered the monocyte-to-neutrophil ratio. In non-exposed WT samples, myeloid cells comprised roughly 70% monocytes and 30% neutrophils, whereas CSE exposure shifted this to an even 50/50 distribution (Figure 6B). In contrast, Tet2-deficient cells displayed an opposite baseline ratio of about 60% neutrophils and 40% monocytes, which CSE further skewed to approximately 75% neutrophils and 25% monocytes (Figure 6B). Overall, EVE and CSE enhanced myelopoiesis in Tet2-deficient cells only, with CSE driving a stronger neutrophil bias. In WT cells, CSE did not change total myeloid frequency but significantly altered lineage composition, indicating that CSE affects both genotypes, whereas EVE selectively impacts Tet2-deficient cells.

### CSE and EVE increase the proportion of Tet2-deficient inflammatory Ly6C++hi monocytes

To further assess how smoke exposure influences myeloid cell function, we analyzed the distribution of monocyte subsets in the transplanted mice. Monocytes are functionally heterogeneous and are typically classified based on their surface marker expression. In mice, subsets are defined by Ly6C expression: Ly6C++high monocytes correspond to classical “inflammatory” monocytes (McGrath et al. 2015; Trzebanski and Jung 2020; Li et al. 2022), which rapidly respond to immune challenges and exhibit strong phagocytic and inflammatory capacity. Ly6C-low monocytes represent the nonclassical “patrolling” subset, which contributes to vascular homeostasis, wound healing, and tissue repair (Lessard et al. 2017; Robinson et al. 2021; Wong et al. 2011). The Ly6C+intermediate population has characteristics that overlap with the inflammatory phenotype of the classical subset (Wong et al. 2012). Peripheral blood monocytes of mice transplanted with non-exposed BM were composed of about 20% classical Ly6C++high monocytes, 40% nonclassical Ly6C-low monocytes, and 40% Ly6C+intermediate monocytes (Figure 7A). Mice transplanted with EVE- or CSE-exposed BM displayed a modest increase in inflammatory Ly6C++high monocytes, at the expense of the Ly6C-low subset (Figure 7A). To assess whether these effects differed between genotypes, we examined each subset separately, comparing wild-type (CD45.1^+^) and Tet2^⁻/⁻^ (CD45.1/2^+^) donor cells (Figure 7B). The Tet2^⁻/⁻^ monocytes have a higher relative proportion of Ly6C++hi monocytes than the WT monocytes (Figure 7B). Notably, the mice transplanted with HSCs exposed to EVE, CSE, or CSE+EVE had significantly higher numbers of inflammatory Ly6C++hi monocytes, but selectively in the Tet2^⁻/⁻^ population (Figure 7B).

**Figure 7.**
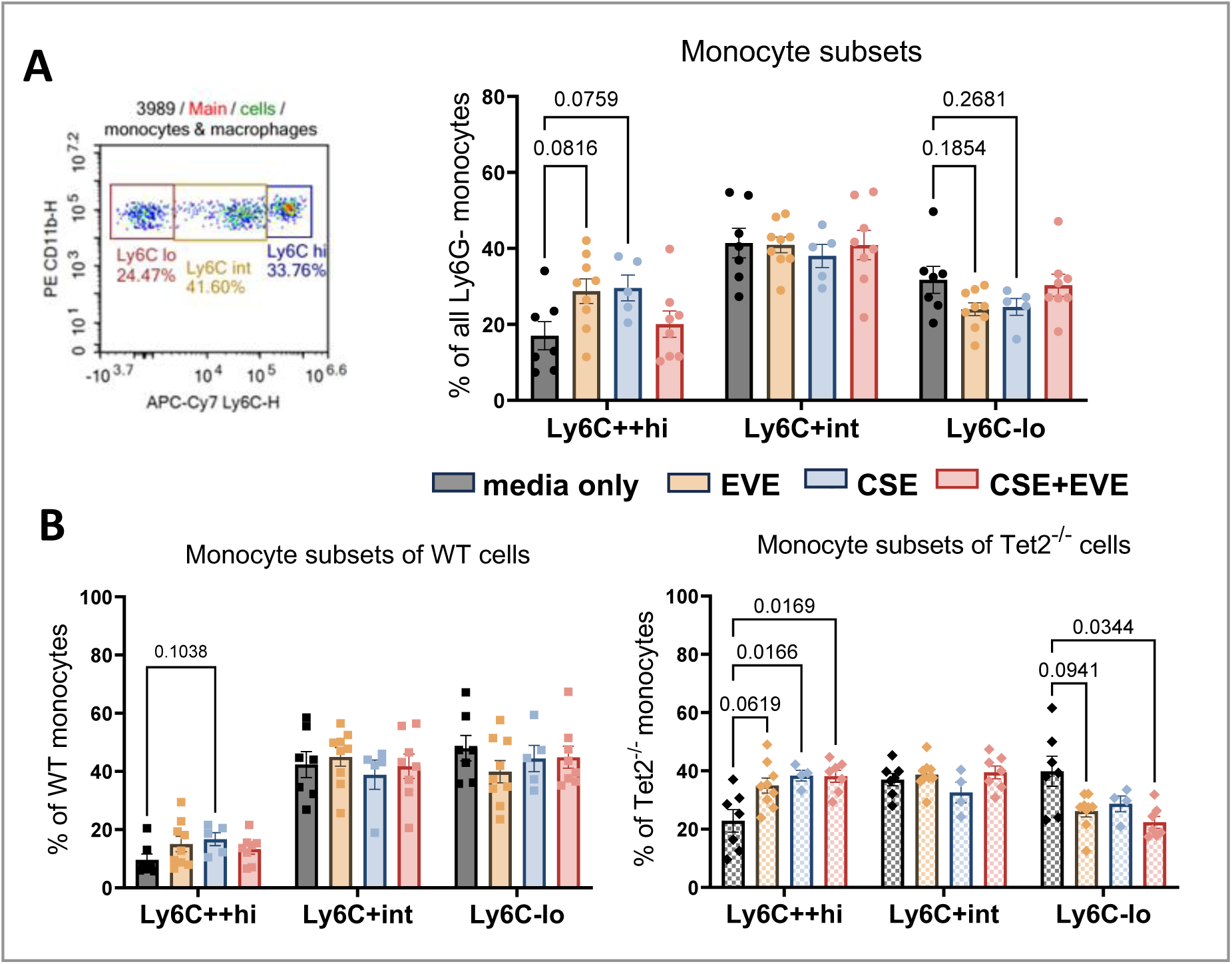
Impact of CSE and EVE on WT and Tet2^-/-^ monocyte subsets. (A) Monocyte subsets of chimeric mice transplanted with CSE- and EVE-exposed bone marrow as defined by relative Ly6C expression measured by flow cytometry. (B) Ly6C subsets of CD45.1 WT (left) or CD45.1/2+ Tet2-/- (right) CD11b+Ly6G-monocytes. Significance was determined by one-way ANOVA. Values represent means ± SEM.

### EVE promotes Tet2^-/-^ proliferation in LT-HSCs in the bone marrow niche

After ten months post-transplant, bone marrow and peripheral blood were collected from the transplanted mice and stained for mature and immature stem/progenitor cell populations. Hematopoietic stem cell populations were identified by staining of the signaling lymphocyte activation molecule (SLAM) family markers CD48 and CD150, with CD48-CD150+ identifying the most primitive long-term HSCs (LT-HSCs, also referred to as LSK-SLAM) (Figure 8A). We observed a significant increase in Tet2^⁻/⁻^ contribution to LKS-SLAM cells in the EVE group compared to the media alone group, demonstrating that Tet2^-/-^ LT-HSC gained a stronger selective advantage over their WT counterparts in the context of a brief ex vivo exposure to EVE (Figure 8B). This expansion of Tet2^⁻/⁻^ cells in the EVE-exposed transplant was also reflected in the peripheral blood throughout the myeloid compartment, with EVE driving proliferation of Tet2-deficient cells through the monocyte (Ly6G-) and neutrophil (Ly6G+) cell types (Figure 8C). In addition to staining for chimerism, we analyzed the frequencies of mature myeloid cells and myeloid progenitors in the bone marrow. EVE slightly increased the frequency of myeloid (CD11b+) cells within the bone marrow and significantly increased the neutrophil bias of these myeloid cells (Supplemental Figure 2A, B).

**Figure 8.**
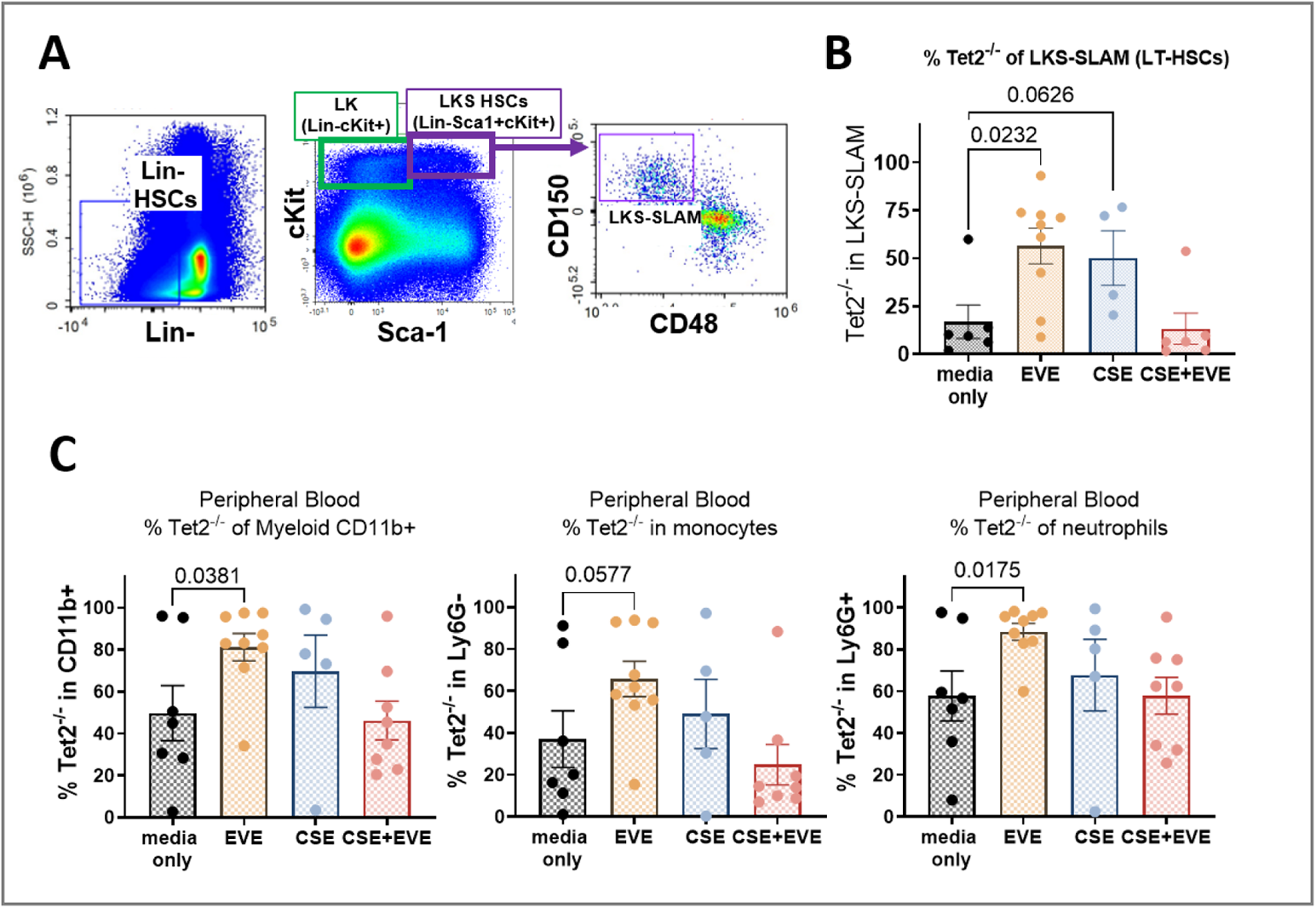
Tet2-/- chimerism in bone marrow at ten months post-competitive transplant with CSE and EVE-exposed HSCs. (A) Representative gating of HSCs. (B) Contribution of Tet2-/- cells to the LKS-SLAM (long-term HSCs) HSCs compartment at termination. (C) Contribution of Tet2-/- cells of the peripheral blood myeloid compartments at termination. Significance was determined by one-way ANOVA. Values represent means ± SEM.

**Figure 9.**
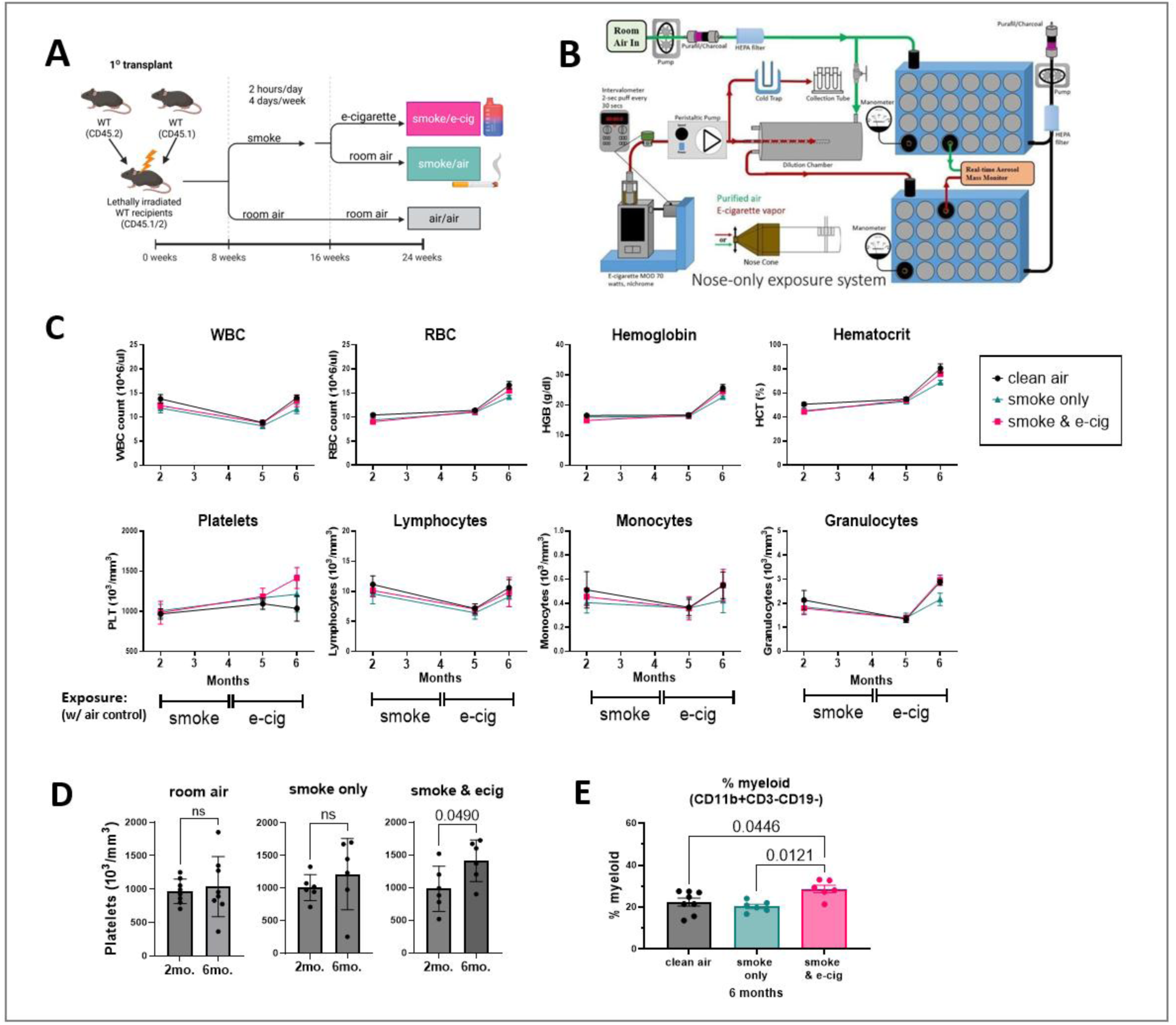
Long-term inhalation of traditional and e-cigarette vape smoke increases the frequency of platelets and myeloid cells in the peripheral blood. (A) Experimental setup of BM transplant and 6-month *in vivo* smoke and e-cigarette exposure. (B) Schematic of custom nose-only exposure system. (C) Complete blood counts of mice during exposure period. (D) Platelet counts at the beginning and end of exposure period. (E) Frequency of CD11b+ myeloid cells of all CD45+ leukocytes in the peripheral blood at end of exposure. Values represent means ± SEM. Significance calculated by paired t-tests (D) and one-way ANOVA (E).

### Cigarette smoke and e-cigarette vape consumption increase the frequency of myeloid cells in the peripheral blood

To model the effects of transitioning from combustible smoking to vaping on hematopoiesis, we exposed mice to sequential combustible smoke and e-cigarette aerosol (Figure 9). A nose-only exposure system was used to ensure uniform delivery and to avoid variability from ingestion or dermal absorption inherent to whole-body exposure (Figure 9A). Mice were exposed to conventional cigarette smoke or clean air for 2 hours per day, 4 days per week (Figure 9B). After two months of smoke exposure, mice in the smoke group were switched to either e-cigarette vapor (n = 6) or clean air (n = 6) (Figure 9B).

Complete blood counts remained largely stable during the exposure period, except for a significant rise in platelet counts in the smoke followed by e-cigarette group (Figure 9D). The most pronounced increase occurred after the switch to e-cigarette exposure (Figure 9C–D), suggesting persistent inflammatory activation following prior smoke exposure. In addition to blood counts, we monitored myeloid and lymphoid cell frequencies in peripheral blood throughout the exposure period and found a myeloid skewing in the smoke followed by e-cigarette group (Figure 9E). At study termination, phagocytic activity of peritoneal macrophages did not differ between groups (Supplemental Figure 3), and the frequency of long-term hematopoietic stem cells in bone marrow was unchanged by either smoke or e-cigarette exposure (Supplemental Figure 3).

### Smoke exposure induces long-lasting myeloid skewing in secondary transplants

To test how smoke or e-cigarette exposure influences the long-term reconstitution capacity of hematopoietic stem cells, we performed secondary bone marrow transplants with bone marrow from the “clean air”, “smoke only”, or “smoke & e-cig” exposed mice (n = 10–13 per exposure group) (Figure 10). Both smoke-only and combined smoke & e-cigarette exposure led to increased frequencies of monocytes and macrophages (CD45⁺CD11b⁺Ly6G⁻) in the peripheral blood with age (Figure 10), consistent with a myeloid bias. Myeloid skewing is a hallmark of hematopoietic aging, and these results suggest that smoke and e-cigarette exposure imprint long-lasting changes on stem cells, which persist through serial transplantation and functionally resemble accelerated HSC aging.

**Figure 10.**
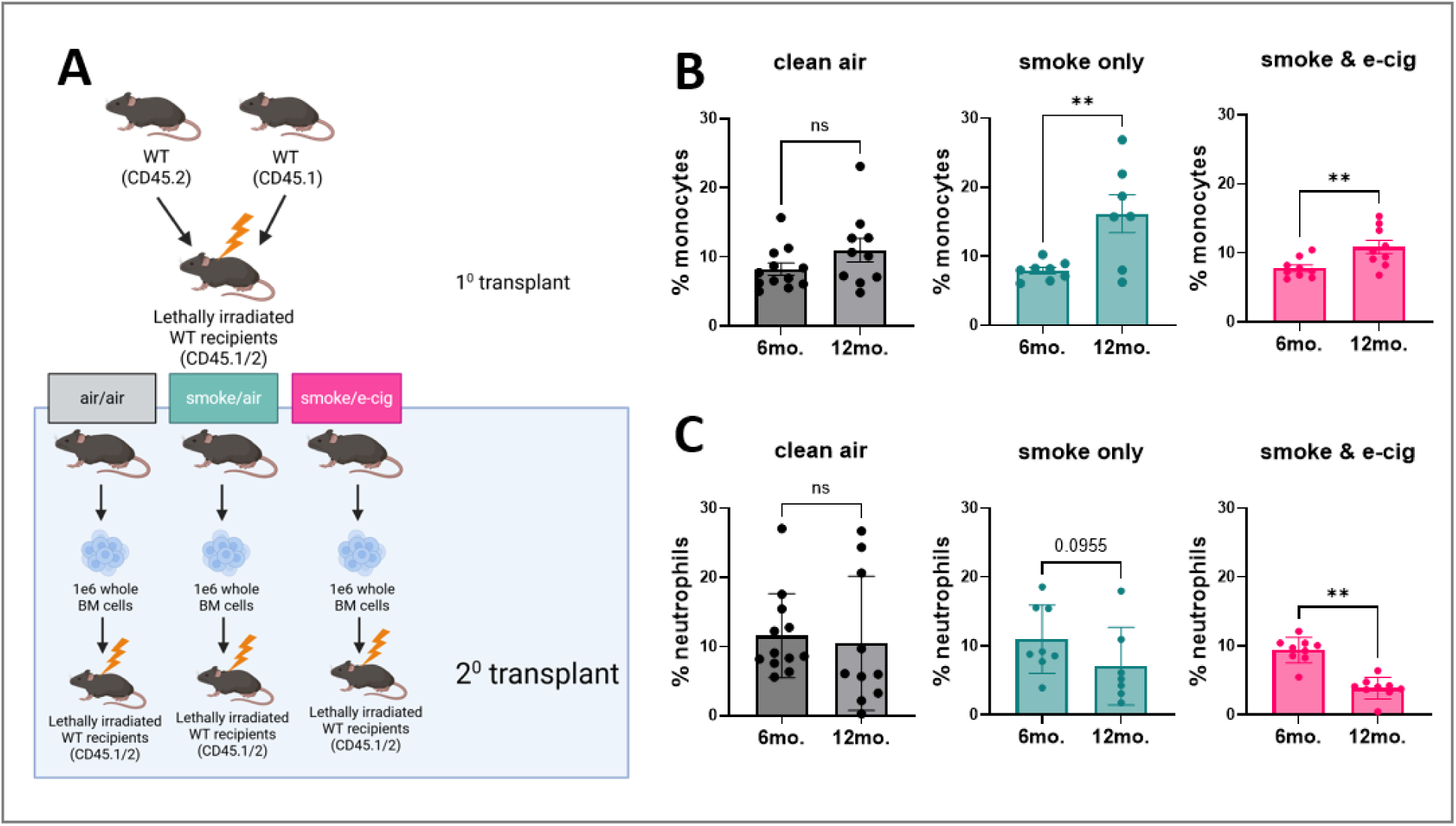
Exposure to cigarette smoke and e-cigarette vapor increases monocytosis and decreases neutrophil differentiation in a secondary transplant. (A) Setup of BM transplant from smoke and e-cigarette-exposed mice into non-exposed mice. (B) Frequency of CD11b+Ly6G- monocytes of all CD45+ leukocytes in the peripheral blood. (C) Frequency of CD11b+Ly6G+ neutrophils of all CD45+ leukocytes in the peripheral blood. Values represent means ± SEM. Significance calculated by paired t-tests.

## DISCUSSION

In this study, we examined the impact of both *in vivo* cigarette smoke and e-cigarette vape inhalation, as well as *in vitro* exposure to concentrated condensed smoke or vape extracts. Evidence suggests that e-cigarette use impacts hematopoiesis and immune cell function (Marques et al. 2021; Scott et al. 2018; Kelesidis et al. 2020). However, the literature is limited and contradictory, and there is very little work on e-cigarettes and clonal hematopoiesis specifically. Extending this investigation from primitive HSCs through the mature myeloid compartment provides insight into how e-cigarettes impact chronic inflammatory responses that can also contribute to mutant hematopoiesis. Here, we found that cigarettes and e-cigarettes increase myelopoiesis in mice and blunt TNF-α secretion in in vitro models of acute immune stimulation.

We first established the impacts of our prepared cigarette smoke extract (CSE) and e-cigarette vape extract (EVE) on mature myeloid cells and HSCs in vitro. Lipopolysaccharide (LPS), or endotoxin, is a component of the outer cell membrane of Gram-negative bacteria and is a potent activator of inflammatory responses. Based on the literature documenting increased levels of inflammatory cytokines in smokers, we anticipated increased TNF-α production in response to CSE or EVE exposure. Contrary to this hypothesis, we observed that CSE suppressed LPS-induced TNF-α secretion. We demonstrated a dose-dependent effect of CSE on human and mouse cell lines, on freshly isolated bone marrow-derived macrophages from wild-type and Tet2^-/-^ mice, and in human PBMCs. This finding encapsulates one of the central debates in nicotine research. Given the complexity of the immune system, it is perhaps unsurprising that there is evidence for both suppression (Ween et al. 2017; Hamon et al. 2024; Ouyang et al. 2000; Arimilli et al. 2017; Breivik et al. 2009; Chen et al., n.d.; Gaschler et al. 2008) and exacerbation (Zhao et al. 2023; Giebe et al. 2023; Mehta and Dhawan 2020) of immune responses by cigarettes and nicotine, depending on the cell and context. Although the finding that CSE inhibits TNF-α seems paradoxical to the increased levels of inflammatory cytokines in smokers, our findings support the paradigm that chronic inflammation impairs an acute immune response. We posit that chronic nicotine use over time increases overall levels of circulating inflammatory cytokines but inhibits the ability to respond to an immune stimulus, hence serving both pro- and anti-inflammatory roles. This is supported by studies such as Elisia et al, who observed that whole blood from smokers has elevated numbers of monocytes and granulocytes and overall increased levels of pro-inflammatory cytokines, but an impaired immune response to an ex vivo acute immune challenge (Elisia et al. 2020). The suppression of inflammatory cytokine production may be a pathway by which cigarette smoke impairs acute immune responses.

We then sought to understand how these smoke-impaired immune responses impact the development of clonal hematopoiesis. Colony formation unit assays are the gold standard for assessing HSC function (Sarma et al. 2010). A fixed number of input hematopoietic progenitors is cultured in a semisolid medium, and the quantity and morphology of colonies formed represent HSC differentiation and proliferation ability. We demonstrated that both CSE and EVE significantly impair HSC colony formation in both WT and Tet2-deficient cells, however Tet2^-/-^ are less impacted compared to WT HSC. El-Mouelhy et al found significantly reduced CFU after exposure to cigarette smoke but not a milk and honey e-cigarette (El-Mouelhy et al. 2022). Our results are consistent with the literature on traditional combustible tobacco cigarettes and reinforce the heterogeneity of e-cigarette products and their impacts.

Studies of the health impacts of cigarette and e-cigarette smoking have naturally been primarily focused on the lungs (Seiler et al. 2020). Smoking has been shown to increase the severity of COPD (Kastratovic et al. 2024; Han et al. 2021; Gong et al. 2019). The lungs contain abundant immune cells, especially macrophages, which originate from both embryonic and hematopoietic stem cell origins (Wohnhaas et al. 2024). Interestingly, individuals with CHIP have both an increased risk of COPD and increased disease severity (Miller et al. 2022). In fact, Miller et al. demonstrated that mice with hematopoietic Tet2 deficiency developed significantly more emphysema after 6 months of cigarette smoke exposure, notably without expansion of the Tet2^-/-^ cells. We have previously shown that 2 months of nose-only exposure to traditional combustible smoke promotes the expansion of Tet2^-/-^ cells (Ramanathan et al. 2023). In this study, we combine in vitro CSE and EVE exposures with in vivo transplants to elucidate a novel dynamic of how smoking impacts hematopoiesis. To our knowledge, this is the first study that tracks the differentiation dynamics of HSCs exposed to cigarette and e-cigarette extracts in an in vivo bone marrow transplant. Competitive BM transplants are a well-established method of assessing relative proliferation and fitness between wild-type and mutant HSCs. Using this competitive model, we demonstrate that exposure to e-cigarette vape extract provides a selective advantage to Tet2-deficient cells. Mice transplanted with EVE-exposed BM had increased frequencies of Tet2-deficient cells in the bone marrow and peripheral blood several months post-transplant. Additionally, mice transplanted with both EVE- and/or CSE-exposed BM had deranged hematopoiesis at over 9 months post-exposure, demonstrating that smoke products have enduring impacts on HSCs. The disparity in the neutrophil-to-monocyte ratio observed between Tet2-/- and WT cells indicates that, in a chimeric model, Tet2-/- myeloid progenitors favor neutrophil differentiation over monocyte differentiation. In contrast, the WT cells do the opposite. Cigarette smoke increased neutrophil frequencies in both WT and Tet2-deficient cells.

We identified that both CSE and EVE increase the frequency of inflammatory (Ly6C++hi) classical monocytes. Characterizing monocyte heterogeneity has redefined our understanding of these cells and identified new potential mechanisms for controlling inflammation. Human *in vivo* deuterium labeling suggests that classical monocytes leave the bone marrow first and circulate for about a day, while intermediate and nonclassical monocytes have slightly longer lifespans of ∼4 and ∼7 days, respectively (Patel et al. 2017). The quantity and relative frequencies are altered under pathological stressors such as infection, cancer, cardiac events, and inflammatory or autoimmune diseases (Kratofil et al. 2017). Monocyte subsets distributions differ in the peripheral blood, BM, lymph nodes, and spleen, depending on local needs (Ożańska et al. 2020). Idel et al observed that smokers have significantly increased expression of the integrin CD11b, an immune response suppressor, on their classical and intermediate monocyte subsets (Idel et al. 2022). Manipulating monocyte subsets to promote patrolling monocytes over inflammatory classical monocytes is being explored as a potential therapeutic approach for autoimmune diseases.

To extend our study, we performed long-term nose-only inhalation exposures to traditional combustible cigarette smoke followed by a switch to either room air (smoking cessation) or to e-cigarette aerosols (switching from smoking to vaping). Our findings are consistent with Verma et al., who observed increased myeloid progenitors and decreased colony-forming ability in mice exposed to cigarette smoke 3 days/week for 4 weeks (Verma et al. 2025). A key distinction between our in vitro CSE and EVE cell exposures and the nose-only inhalations is that the EVE is created using a commercially available melon-flavored e-liquid. In contrast, to eliminate commercial manufacturing variables, the inhaled e-cigarette aerosol is generated from house-made e-liquid containing a 50/50 PG/VG ratio and 1.5% nicotine. Including both commercial and house-made e-cigarette formulations in these studies highlights that even the base components alone alter hematopoiesis.

These long-term cellular analyses of the impact of e-cigarettes or traditional tobacco cigarettes on hematopoiesis provide insight into how smoking history may impact the progression of CHIP. We conclude that both combustible and electronic cigarette smoking derange hematopoiesis to distort myeloid differentiation. Importantly, these results suggest that the impacts of even minimal exposure to smoke extracts are persistent after smoking cessation. Although we can tell smokers to cease, we cannot erase their smoking history. Therefore, understanding how smoking impacts the body, even after cessation, is of critical importance when identifying effective therapeutics. As the population ages, incorporating studies of clonal hematopoiesis in disease contexts is of increasing clinical concern. Our results demonstrate that exposure to both cigarettes and e-cigarettes permanently changes long-term HSC myeloid lineage decisions.

## Author contributions

JYCS designed and performed the experiments, analyzed the data, and wrote the manuscript; HXH, CB, FD, and JHC performed experiments; NI and NLS provided reagents; DH, IH, AT, and MK developed and oversaw or performed nose-only exposures; AGF developed the experimental plan, oversaw research, analyzed data, and edited the manuscript.

## Acknowledgements

We thank Irene Hasen and Amanda Ting in the Air Pollution and Health Effects Laboratory at the University of California, Irvine, for their assistance with the in vivo exposures.

