## Supplemental Figures for "Cigarette Smoke and E-Cigarette Aerosol Extracts Induce Myelopoiesis and Suppress Inflammatory Cytokine Production"

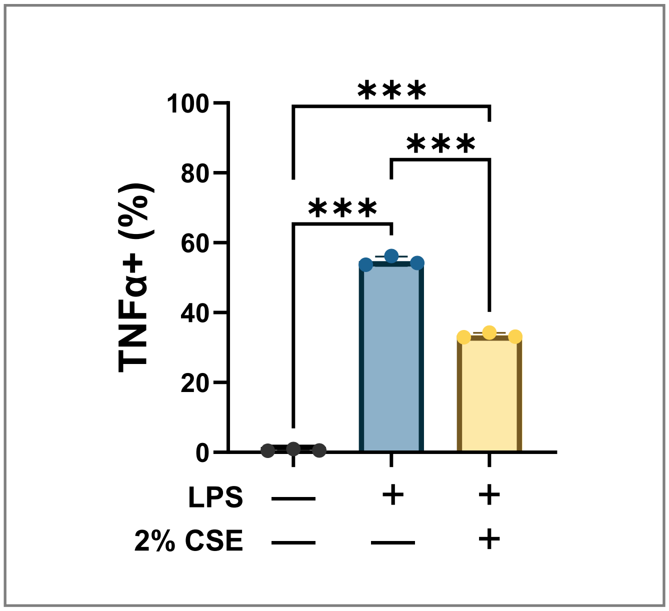


**Supplemental Figure 1. Cigarette Smoke Extract suppresses LPS induced TNF-α production.** RAW264.7 mouse macrophages were stimulated with LPS (10ng/mL) +/- 2% CSE for 8 hours, then harvested, fixed, permeabilized and stained with TNF-α antibody to detect the % of TNF-α+ cells. Significance is calculated by one-way ANOVA. Values represent means ± SEM.

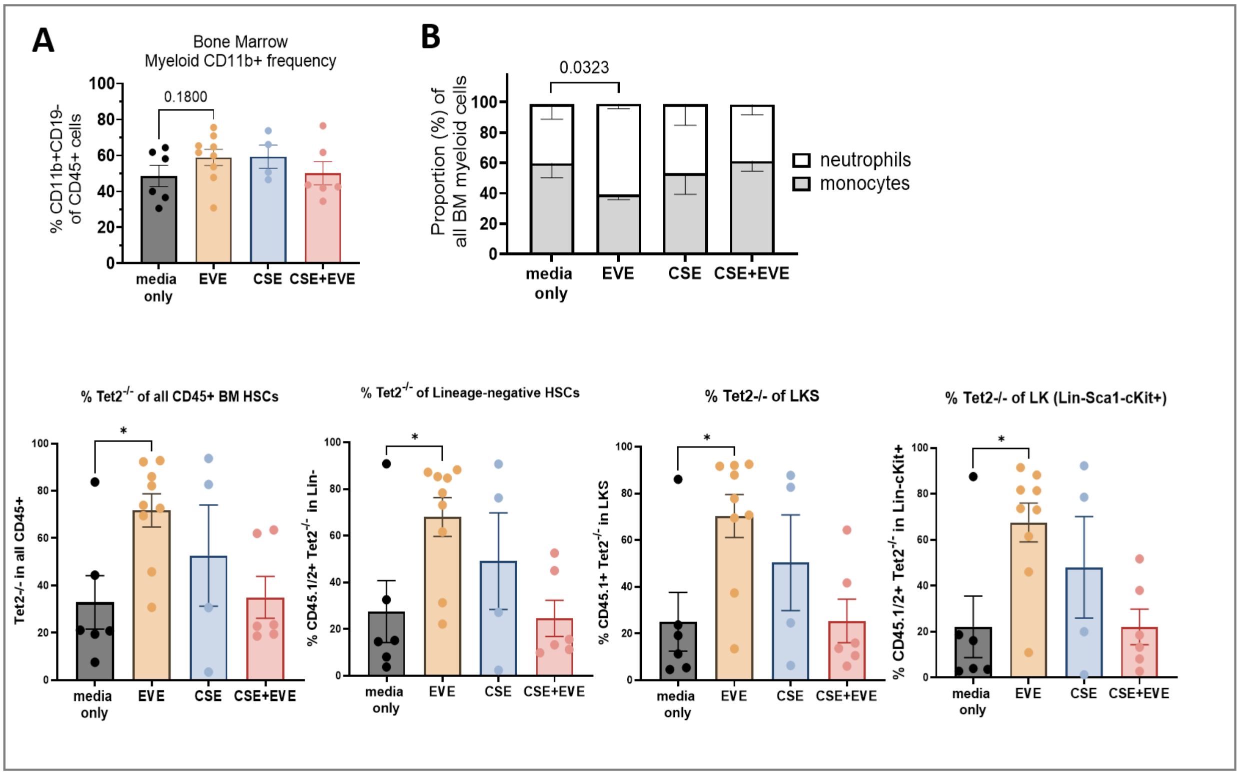


**Supplemental Figure 2. Tet2 Analysis of Tet2-/- and myeloid cell frequencies in the bone marrow at ten months post-competitive transplant with CSE and EVE-exposed HSCs.** (A) Frequency of CD11b+ myeloid cells of all CD45+ bone marrow. (B) Proportions of Ly6G+ neutrophils and Ly6G- monocytes in myeloid bone marrow cells (CD45+CD11b+). (C) Frequency of Tet2-knockout cells in hematopoietic stem cell populations. Significance is calculated by one-way ANOVA. Values represent means ± SEM.

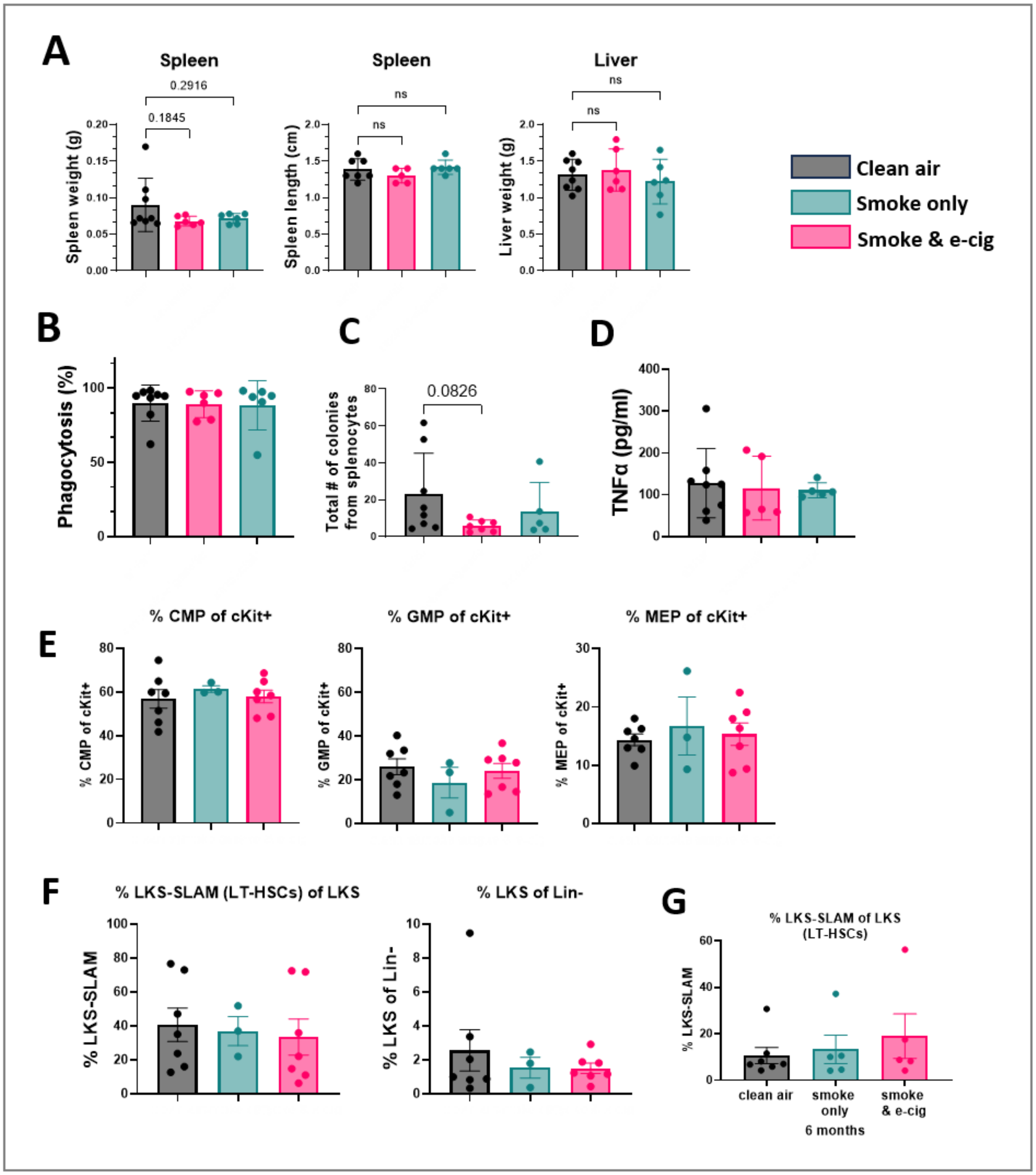


**Supplemental Figure 3. In vivo smoke exposure termination analyses.** (A) spleen weight and length, and liver weight, analyzed by one-way ANOVA. (B) Peritoneal macrophage phagocytosis. Peritoneal macrophages were assessed for phagocytic capacity by incubation with fluorescent Zymosan beads. (C) Roughly half of the spleen was crushed and cells harvested through a 0.22uM filter. For colony formation, 100k splenocytes were plated into 1mL methylcellulose + mSCF and mIL-3 and counted 7 days later. (D) Splenocytes were plated and challenged with 100ng/mL LPS for 24 hours. Wells were collected and spun down and supernatant was removed from the cell pellet and frozen at -80 until analysis. (E) CMP, MEP, GMP of cKit+ HSCs from exposed mice. (F) LKS and LKS-SLAM analysis of HSCs from exposed mice. (G) LKS-SLAM analysis of HSCs from mice transplanted with BM from smoke and e-cig-exposed mice.
